# Lineage-specific chromatin poising enforced by ETS-IRF composite elements determines the divergent interferon responses of plasmacytoid dendritic cells and epithelial cells

**DOI:** 10.64898/2026.06.05.730516

**Authors:** Gargi Mishra, Snehal Ozarkar, Yo Okamura, Meghan Knoll, Russell Sam, Hongchuan Li, Johannes Schwerk, Adelle P. McFarland, David Constant, Hayley Waterman, Manasa Acharya, Adam Lacy-Hulbert, Jason G. Smith, Michael Parthun, Emily Hemann, Isaac M Barber-Axthelm, Weiming Li, Vishal Nigam, Timothy Nice, Jessica A. Hammerman, Stephen K Anderson, Ram Savan

**Affiliations:** Department of Immunology, School of Medicine, University of Washington, Seattle, WA, USA; Basic Science Program, Frederick National Laboratory for Cancer Research, Frederick, MD, USA; Oregon Health & Science University, Portland, Oregon, USA; Centre for Fundamental Immunology, Benaroya Research Institute, Seattle, WA, USA; Department of Microbiology, University of Washington, Seattle, WA, USA; Department of Microbial Infection and Immunity, The Ohio State University College of Medicine, Columbus, Ohio, USA; Department of Comparative Medicine, School of Medicine, University of Washington, Seattle, WA, USA; Department of Pediatrics (Cardiology), University of Washington, Seattle, WA, USA; Center for Developmental Biology and Regenerative Medicine, Seattle Children’s Research Institute, Seattle, WA, USA; Cancer Innovation Laboratory, Center for Cancer Research, NCI, Frederick, MD, USA

## Abstract

Plasmacytoid dendritic cells (pDCs) produce robust type I interferons (IFN-I) within hours of viral sensing, while epithelial cells at mucosal surfaces mount a delayed response dominated by type III interferons (IFN-III). Both cell types express pattern recognition receptors that activate similar downstream transcription factors, yet they produce distinct subsets of IFNs. The mechanisms underlying these differences have remained unclear. Here, using Assay for Transposase-Accessible Chromatin using sequencing (ATAC-seq) in primary human pDCs and intestinal epithelial cells, we show that IFN-I and IFN-III gene loci carry opposing, constitutively established chromatin accessibility landscapes that determine cell-type-specific interferon induction. The IFN-I locus is broadly open in pDCs and constitutively closed in epithelial cells, while the IFN-III locus displays the reciprocal pattern. Motif enrichment analysis of accessible regions at the IFN-I locus in pDCs revealed unexpected and significant enrichment of ETS family binding motifs alongside IRF motifs, which determines the cell-type-specific locus accessibility. The ETS factor PU.1 and IRF8 bind IFN-I promoters, at composite ETS-IRF elements positioned at the Positive Regulatory Domain IV (PRDIV) site of *IFNB1* promoter and adjacent to the TATA-proximal IRF motif of IFNA promoters. *IFNL* gene promoters lack ETS recognition sequences. PU.1-IRF8 composite factor binding extends across intergenic regions of the IFN-I locus, where candidate enhancer elements marked by H3K4me1, H3K27ac, and RNA Pol II occupancy were identified. We propose that this network of ETS-IRF composite-element-anchored enhancers maintains the IFN-I locus in a constitutively poised state in pDCs, licensing the rapid and robust IFN-I response that defines pDCs. In epithelial cells, the absence of PU.1 and IRF8 renders the IFN-I locus epigenetically silent, while the IFN-III locus is constitutively open. Despite this, the delayed IFN-III gene expression in epithelial cells is due to intrinsically weaker promoter activity relative to IFN-I. These findings reveal that the divergent IFN induction of pDCs and epithelial cells is determined by the chromatin architecture prior to infection. Overall, these observations show that lineage-specific ETS-IRF regulatory factors and promoter strength determine the cell-type-specific IFN activation.

## Introduction

Type I and III interferons (IFNs) are critical antiviral effectors essential for virus clearance^1–3^. Type I interferons (IFN-I; α/β/m/E/K) are considered the dominant interferons that contribute to priming both innate and adaptive arms of the immune system to protect against all virus infections^4^. IFN-I are required for the induction of a robust adaptive antiviral immune response critical for sterilizing immunity. They also trigger apoptosis of infected cells to halt viral replication & induce an antiviral state in neighboring uninfected cells through paracrine signaling, thereby containing viral dissemination. IFN-III (λ) action is more localized to mucosal tissues and primarily establishes an innate antiviral state by inducing antiviral restriction factors^5,6^. IFN-III is critical for clearing mucosal pathogens such as hepatitis C virus (HCV)^7–14^, norovirus^15–17^, rhinovirus, respiratory syncytial virus, influenza A virus^18^, hepatitis B virus, rotavirus, and reovirus [reviewed in ^5,19–22^]. These observations suggest non-redundant but collaborative actions of IFN-I and IFN-III.

Sensing and induction pathways leading to IFN production are critical for pathogen clearance as well as for interferonopathies^23^. IFN-I & IFN-III are induced upon recognition of pathogen-associated molecular patterns (PAMPs) derived from viral replication intermediates or self-nucleic acids through pathogen recognition receptors (PRRs). These include toll-like receptors (TLRs)^24–27^, RNA sensors such as RIG-I-like receptors (RLR; RIG-I and MDA5)^28–31^, and DNA sensors including cGAS^32^ and IFI16^33^. Engagement of these receptors initiates a signaling cascade that culminates in the activation of IRF3, IRF5, or IRF7, together with components of the NF-ΚB^34,35^. These transcription factors bind to the promoters of IFN-I genes (comprising 13 *IFNA* subtypes, *IFNB1*, *IFNW1*, and *IFNE*^36,37^) and IFN-III genes (*IFNL1*–*IFNL4*^2,3,14,38^) to initiate their transcription. Despite pattern recognition receptors activating similar transcription factors that bind to comparable promoter motifs, IFN-I and IFN-III genes are cell-type-specific and follow a temporally distinct pattern. IFN-I production is dominated by plasmacytoid dendritic cells (pDCs), which uniquely mount a rapid and robust response early during infection that resolves within hours. While not produced in blood-derived pDCs, IFN-III is primarily produced by epithelial cells at mucosal surfaces, the critical sites of entry for most viruses^39^. Unlike the transient yet rapid IFN-I burst from pDCs, epithelial IFN-I and IFN-III induction is delayed. The molecular basis for these temporal and cell-type-specific differences in IFN gene induction has remained unclear. In the present study, we identify novel cell-type-specific regulatory mechanisms, operating at the level of chromatin accessibility and lineage-restricted transcription factor binding, that govern the distinct IFN-I and IFN-III programs of pDCs and epithelial cells.

## Results

### ATAC-seq reveals a distinct and reciprocal accessibility of IFN-I and IFN-III loci in pDCs and intestinal epithelial cells

Activation of PRRs activate IRF3, IRF5, and IRF7, together with c-JUN/ATF and NF-κB (p50/p65), which cooperate to induce IFN-I and IFN-III gene transcription. Yet the activity of these shared transcription factors alone does not account for the striking cell-type specificity of IFN gene induction: IFN-I production is dominated by pDCs, while epithelial cells at mucosal surfaces are the principal source of IFN-III. We therefore hypothesized that differential chromatin accessibility at IFN gene loci is a primary determinant of the cell-type-specific and temporally distinct induction of IFN-I and IFN-III. To test this, we profiled genome-wide chromatin accessibility by Assay for Transposase-Accessible Chromatin using sequencing (ATAC-seq) in primary human pDCs stimulated with the pathogen-associated molecular patterns (PAMPs) R848, a small molecule TLR7/8 ligand^40,41^, and compared to human intestinal epithelial organoids stimulated with a RIG-I ligand (RIG-IL), alongside matched mock-treated controls. ATAC-seq uses the Tn5 transposase to insert sequencing adapters preferentially into nucleosome-free genomic regions, providing an unbiased, genome-wide map of open chromatin^42^. Sequencing reads were mapped to define regions of accessibility corresponding to transcription factor occupancy and nucleosome positioning. These comparative analyses revealed strikingly different chromatin accessibility at the IFN-I and IFN-III loci in the two cell types (Fig. 1). In pDCs, the IFN-I locus was highly accessible, while the IFN-III locus remained closed (Fig. 1A, B). At the individual promoter level, strong ATAC-seq peaks were detected near the promoters of *IFNA2* and *IFNE* in unstimulated pDCs, with intermediate accessibility at *IFNA14* and *IFNA5*, and low accessibility at *IFNA13* and *IFNW1* (Fig. 1C). By 3 h post R848 stimulation, accessibility increased markedly and extended across the locus, encompassing *IFNA2*, *IFNE*, *IFNA5*, *IFNA14*, *IFNB1*, and *IFNW1*, suggesting activation-dependent chromatin remodeling in an already poised locus. No accessibility was detected proximal to IFNL promoters at any time point in pDCs before or after R848 treatment.

**Figure 1:**
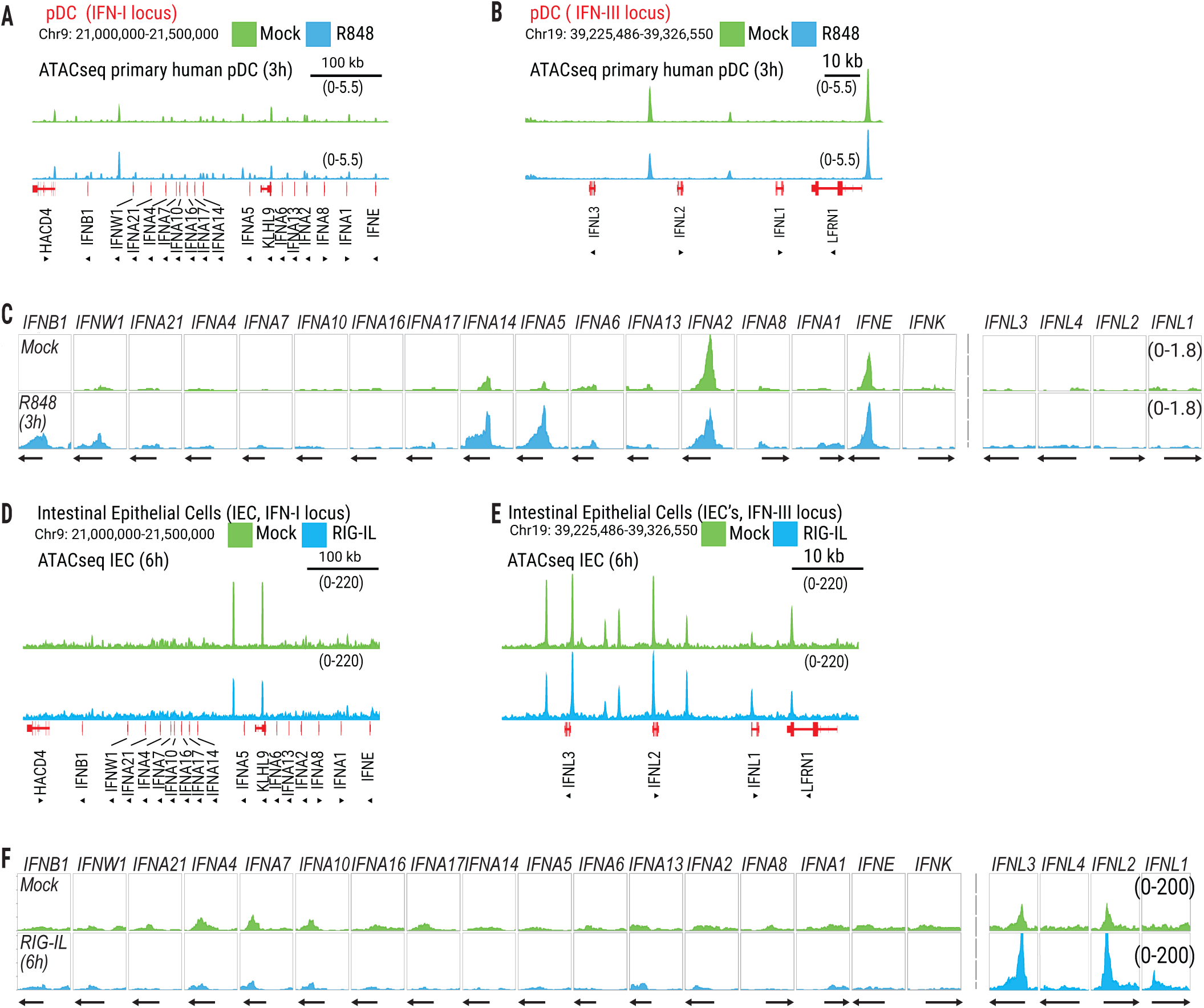
Cell-type-specific chromatin accessibility at the IFN-I and IFN-III loci in human blood plasmacytoid dendritic cells and intestinal epithelial cells. (A-B) Chromatin accessibility of human blood plasmacytoid dendritic cells at the IFN-I locus at 3 h (chr9: 21.0-21.5 Mb) **(A)** and IFN-III locus (chr19: 39,225,486-39,326,550) **(B)**. **(C)** Chromatin accessibility at individual IFN-I and IFN-III gene promoters (1 kb upstream of TSS; gray dashed vertical lines indicate TSS position) in pDCs. **(D-E)** Chromatin accessibility for human intestinal epithelial cells (IECs) derived from intestinal organoids at 6 hr at the IFN-I locus (chr9: 21.0-21.5 Mb) **(D)** and for the IFN-III locus (chr19: 39,225,486-39,326,550) **(E). (F)** Chromatin accessibility at individual IFN-I and IFN-III gene promoters (1 kb upstream of TSS; gray dashed vertical lines indicate TSS position) for IECs. Mock (green) and R848-treated (blue). The data shown are of one representative of 3 biological replicates.

In epithelial organoids, the accessibility pattern was reversed. Under basal conditions, the IFN-III locus was constitutively open, with prominent accessibility peaks at the IFNL2 and IFNL3 gene promoters (Fig. 1D, E). Stimulation with a RIG-I ligand further increased and extended accessibility to include IFNL1. The IFN-I locus was largely inaccessible in epithelial cells (Fig. 1F). Taken together, these results demonstrate that cell-type-specific chromatin accessibility at IFN gene loci is established prior to stimulation and is further remodeled upon PAMP sensing, providing a structural basis for the differential induction of IFN-I in pDCs and IFN-III in epithelial cells. This indicates that the distinct expression patterns of IFN-I and IFN-III genes in epithelial cells versus pDCs are controlled by regional chromatin accessibility and histone marks.

### Cell line models faithfully recapitulate the IFN locus chromatin architecture of their primary cell counterparts

Primary pDCs and intestinal epithelial cells are both scarce and refractory to genetic manipulation, posing significant barriers to understanding the molecular mechanisms of IFN locus regulation. To identify tractable cellular models for follow-up studies, we compared chromatin accessibility profiles at the IFN-I and IFN-III loci between primary cells and their respective cell line counterparts, CAL-1 cell line^43^ for pDCs and the PH5CH8 hepatocyte-derived cell line^44^ for intestinal epithelial cells (IECs). Both CAL-1^45–47^ and PH5CH8^48^ cell lines have been used extensively for the study of interferon and innate immune responses. ATAC-seq was performed under both mock and PAMP-stimulated conditions to assess whether cell line accessibility profiles mimicked those of primary cells across cell states.

In the pDC comparison, accessibility peaks across the IFN-I locus were virtually identical between primary pDCs and CAL-1 cells in both mock and stimulated conditions (Fig. 2A-D). The same promoter-proximal peaks detected in primary pDCs at *IFNA2*, *IFNE*, *IFNA5*, *IFNA14*, *IFNA6*, *IFNB1*, and *IFNW1* were recapitulated in CAL-1 cells, with no substantive differences in peak position (Fig. 1C, 2E). Critically, the IFN-III locus remained inaccessible in CAL-1 cells under all conditions, mirroring the constitutively closed configuration observed in primary pDCs (Fig 1C, 2C-D). Accessibility profiles were also stable between mock and PAMP-stimulated conditions in CAL-1 for up to 24h (Fig. 2C), confirming that the open IFN-I locus architecture in pDC-lineage cells is a constitutive, stimulus-independent feature of this lineage.

**Figure 2:**
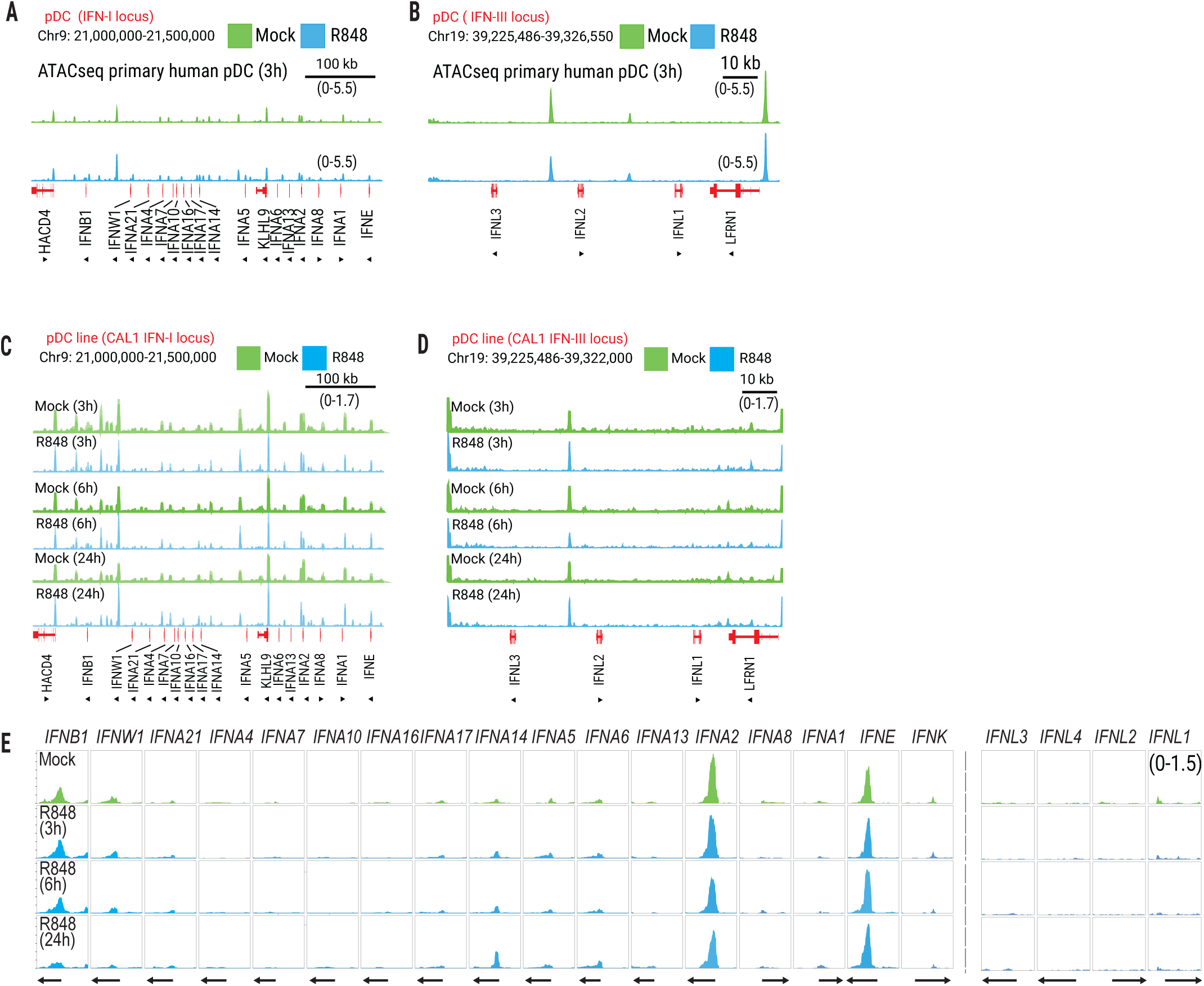
The IFN-I locus is constitutively accessible in naïve and activated CAL-1 cells [pDC cell line] similar to primary human pDCs, while the IFN-III locus remains closed. (A-B) Chromatin accessibility of primary human pDCs at the IFN-I locus primary (chr9: 21.0-21.5 Mb) **(A)** and IFN-III locus (chr19: 39,225,486-39,326,550) **(B). (C-D)** Chromatin accessibility of CAL-1 pDCs at the IFN-I locus primary (chr9: 21.0-21.5 Mb) **(C)** and IFN-III locus (chr19: 39,225,486-39,326,550) **(D)** at the IFN-I locus at 3 h, 6 h, and 24 h. **(E)** Chromatin accessibility at individual IFN-I (left) and IFN-III (right) gene promoters (1 kb upstream of TSS; gray dashed vertical lines indicate TSS position) in CAL-1 cells at 3 h, 6 h, and 24 h. Mock (green) and R848-stimulated (blue). Data in (A-B) are shown from one representative donor. Data in (C-E) are representative of one of 2 biological replicates.

Similar results were observed in the epithelial cell comparison. PH5CH8 cells displayed chromatin accessibility profiles at the IFN-I and IFN-III loci that were largely similar to those of primary IECs under both mock and PAMP-stimulated conditions (Fig. 3A-E). The constitutively open IFN-III locus and the largely inaccessible IFN-I locus characteristic of primary epithelial cells were recapitulated in PH5CH8 cells, with comparable peak distribution and signal across individual *IFNL* promoters in both stimulated and unstimulated states (Fig. 3A-E). PAMP stimulation did not significantly change the chromatin accessibility in either primary IECs or PH5CH8 cells, supporting our conclusion that locus accessibility in epithelial cells is determined by a pre-established chromatin state rather than by stimulus-induced changes.

**Figure 3:**
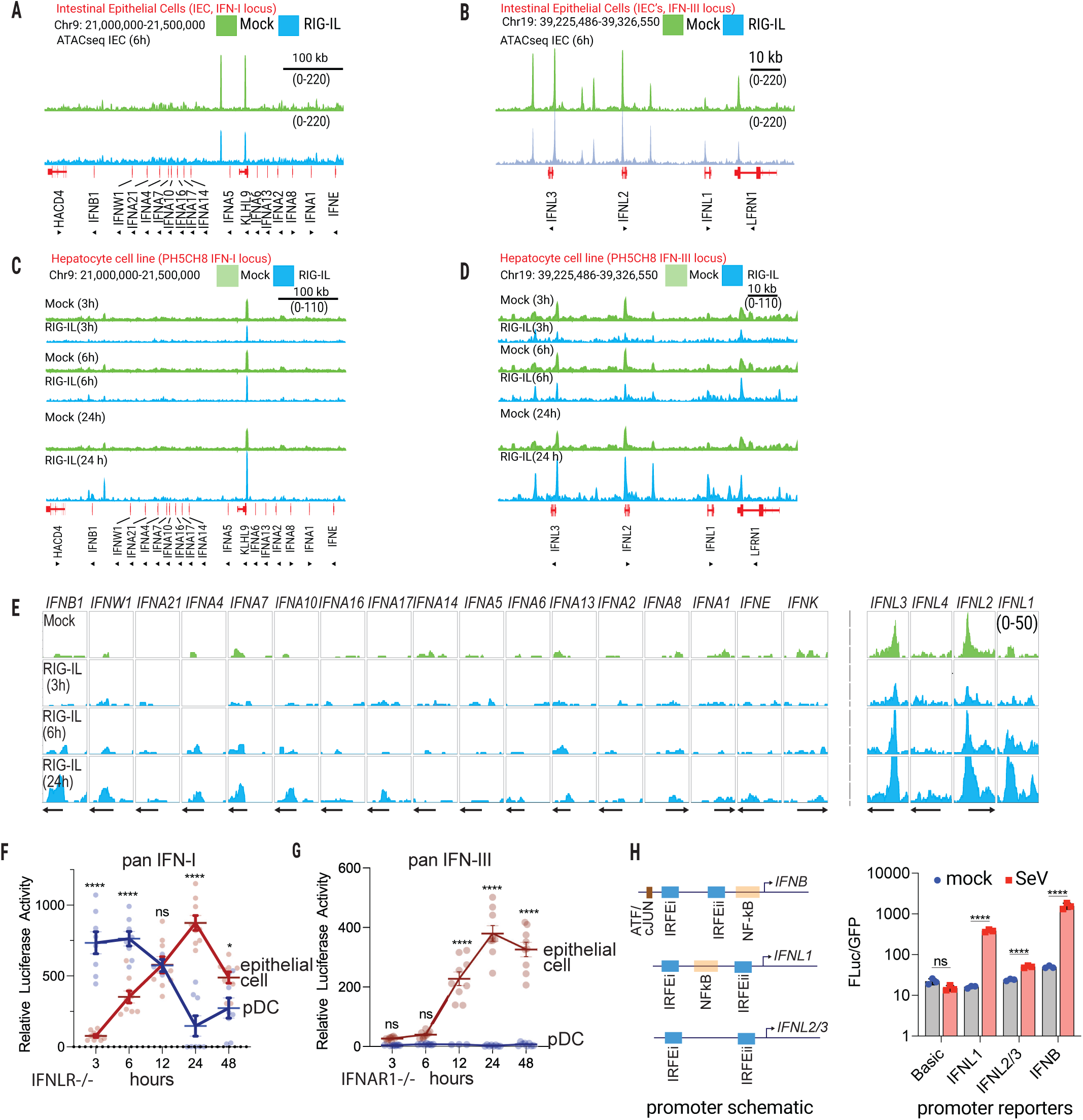
Cell type-specific chromatin accessibility determines IFN-I and IFN-III secretion. (A-B) Chromatin accessibility of primary intestinal epithelial cells (IECs) derived from intestinal organoids at the IFN-I locus (chr9: 21.0-21.5 Mb) **(A)** and the IFN-III locus (chr19: 39,225,486-39,326,550) **(B)** at 6 h. **(C-D)** Chromatin accessibility of PH5CH8 hepatocytes at the IFN-I locus (chr9: 21.0-21.5 Mb) **(C)** and IFN-III locus (chr19: 39,225,486-39,326,550) **(D)** at 3 h, 6 h, and 24 h. **(E)** Chromatin accessibility at individual IFN-I (left) and IFN-III (right) gene promoters (1 kb upstream of TSS; gray dashed vertical lines indicate TSS position) in PH5CH8 hepatocytes at 3 h, 6 h, and 24 h. **(F-G)** Pan IFN-I assay in CAL-1 pDCs and PH5CH8 hepatocytes measured by Gaussia luciferase assay using supernatants transferred to IFNLR1-knockout HuH7 promoter reporter cells **(F)** and IFNAR1-knockout promoter reporter HuH7 cells **(G)** at the indicated time points (0 - 48 h) after PAMP stimulation. **(H) Left**: Schematic of regulatory elements architecture at IFNB1, IFNL1, and IFNL2/3 promoters showing IRFEi, IRFEii, ETS, and NF-ΚB binding elements. **Righ**t: promoter luciferase reporter activity (Fluc/GFP ratio) for IFNB1, IFNL1, and IFNL2/3 and basic promoter reporter constructs in mock and Sendai virus-treated HEK293 cells. Mock (green) and RIG-I ligand-stimulated (blue). Data in **(A-B)** are shown from one representative of two biological replicate. Data in **(C-E)** are representative of one of 2 biological replicates. Data in **(F-G)** are mean ± SEM, n = 3 biological replicates. ***p < 0.001; ns, not significant.

The near-identical chromatin accessibility landscapes between primary cells and their cell line counterparts, across both loci, both cell types and stimulation conditions, establish that CAL-1 and PH5CH8 cells preserve the lineage-specific epigenetic identity of the cell types they model. Because both cell lines are amenable to genetic perturbation, including transcription factor knockdown, overexpression, and CRISPR-based editing, they provide a tractable system for dissecting the mechanisms that influence the chromatin architectures of the IFN-I and IFN-III loci in pDCs and epithelial cells. All subsequent mechanistic analyses were therefore performed in these cell models.

### Cell-type-specific and temporally distinct interferon responses in pDCs and epithelial cells

To measure IFN-I and IFN-III production, we activated CAL-1 and PH5CH8 cells over a time course using R848 or RIG-IL, respectively. Rather than measuring individual IFN subtypes, we developed an IFN-specific bioactivity reporter assay that selectively detects either IFN-I or IFN-III signaling.

Specifically, *IFNAR1*^−/−^ and *IFNLR1*^−/−^ reporter lines, expressing a secreted *Gaussia* luciferase under the control of the *ISG15* promoter, were used to measure panIFN-I or panIFN-III activity, respectively. In CAL-1 cells, IFN-I bioactivity was rapidly and robustly induced following stimulation, with peak activity detected as early as 3 h post-stimulation (Fig. 3F). This response was transient; IFN-I activity declined sharply by 24 h, consistent with the known kinetics of pDC interferon responses in primary cells. In accordance with the closed chromatin configuration of the IFN-III locus in pDC-lineage cells, no IFN-III bioactivity was detected at any time point examined, confirming that pDCs selectively produce IFN-I and are not a source of IFN-III under these conditions (Fig. 3G).

Epithelial cells displayed a significantly different temporal expression pattern. Rather than an early burst, IFN induction was gradual, with both panIFN-I and panIFN-III bioactivity accumulating over time and reaching peak levels at 24 h post-stimulation (Fig. 3F-G). The delayed but sustained induction of IFN-I and IFN-III at later time points contrasts sharply with the rapid IFN-I-dominant transcription of pDCs. These data are consistent with the constitutively open IFN-III locus and closed IFN-I locus we observed in epithelial cells by ATAC-seq.

### The *IFNL* promoter is intrinsically weaker than *IFNB1* and is predominantly driven by IRF-dependent activation

To determine whether differences in intrinsic promoter strength contribute to the distinct expression kinetics of IFN-I and IFN-III genes, we measured the transcriptional activity of the *IFNB1*, *IFNL1*, and *IFNL2/3* promoters in isolation from their broader chromatin context. Wild-type promoter sequences were cloned upstream of a firefly luciferase reporter and transfected into HEK293 cells, along with an eGFP construct for transfection control. A promoter-less luciferase construct (Basic) was included to establish the basal level of reporter activity. Cells were infected with Sendai virus (SeV; 60 HAU/mL) or mock-infected as controls, and luciferase activity was quantified at 24h post-infection (Fig. 3H). SeV infection robustly induced *IFNB1* promoter activity, producing significantly higher luciferase activity than either *IFNL1*or *IFNL2/3* promoters following SeV infection. This shows that the delayed protein production of IFN-III genes relative to *IFNB1* is partly influenced by the strength of the core promoter sequence and is independent of chromatin accessibility. Together, these data suggest that pDCs and epithelial cells mount temporally and qualitatively distinct IFN responses. pDCs produce a rapid and robust IFN-I production, while epithelial cells mount a delayed IFN-III production. There is a high correlation with cell-type-specific gene expression, and chromatin accessibility at IFN-I and IFN-III loci provides direct evidence that locus architecture and promoter strength determine both the identity and timing of IFN responses.

### ETS factors preferentially bind to accessible regions within the IFN-I locus in pDCs

To identify transcription factors associated with accessible regions at the IFN-I locus in pDCs, we performed motif enrichment analysis using HOMER (Supplementary Table 1). Our analysis revealed significant enrichment of sequence recognition motifs for interferon regulatory factors (IRFs) and ETS (E26 transformation-specific) family members. Enrichment of IRF motifs was not surprising, given their well-established role in binding to the IFN-I and IFN-III gene promoters required to initiate IFN transcription. In contrast, significant enrichment of ETS factor recognition sequences was unexpected. Among ETS family members^49^, PU.1 is a lineage-defining transcription factor essential for pDC development and function^50,51^, prompting us to investigate whether PU.1 directly associates with the IFN loci. To test whether PU.1 preferentially binds the IFN-I rather than the IFN-III locus, we performed CUT&RUN profiling for PU.1 in mock-and R848-treated CAL-1 cells. This analysis revealed robust PU.1 deposition across the IFN-I locus, whereas no PU.1 deposition was found at the IFN-III locus under mock or stim conditions (Fig. 4A-B). Strikingly, PU.1 deposition across the IFN-I locus superimposed with ATAC-seq defined accessible regions, indicating that PU.1 preferentially occupies accessible chromatin at the IFN-I locus (Supplementary Table 2). These findings suggested that PU.1 is a critical lineage-specific factor associated with the establishment or maintenance of poised or open chromatin across the IFN-I locus in pDCs.

**Figure 4:**
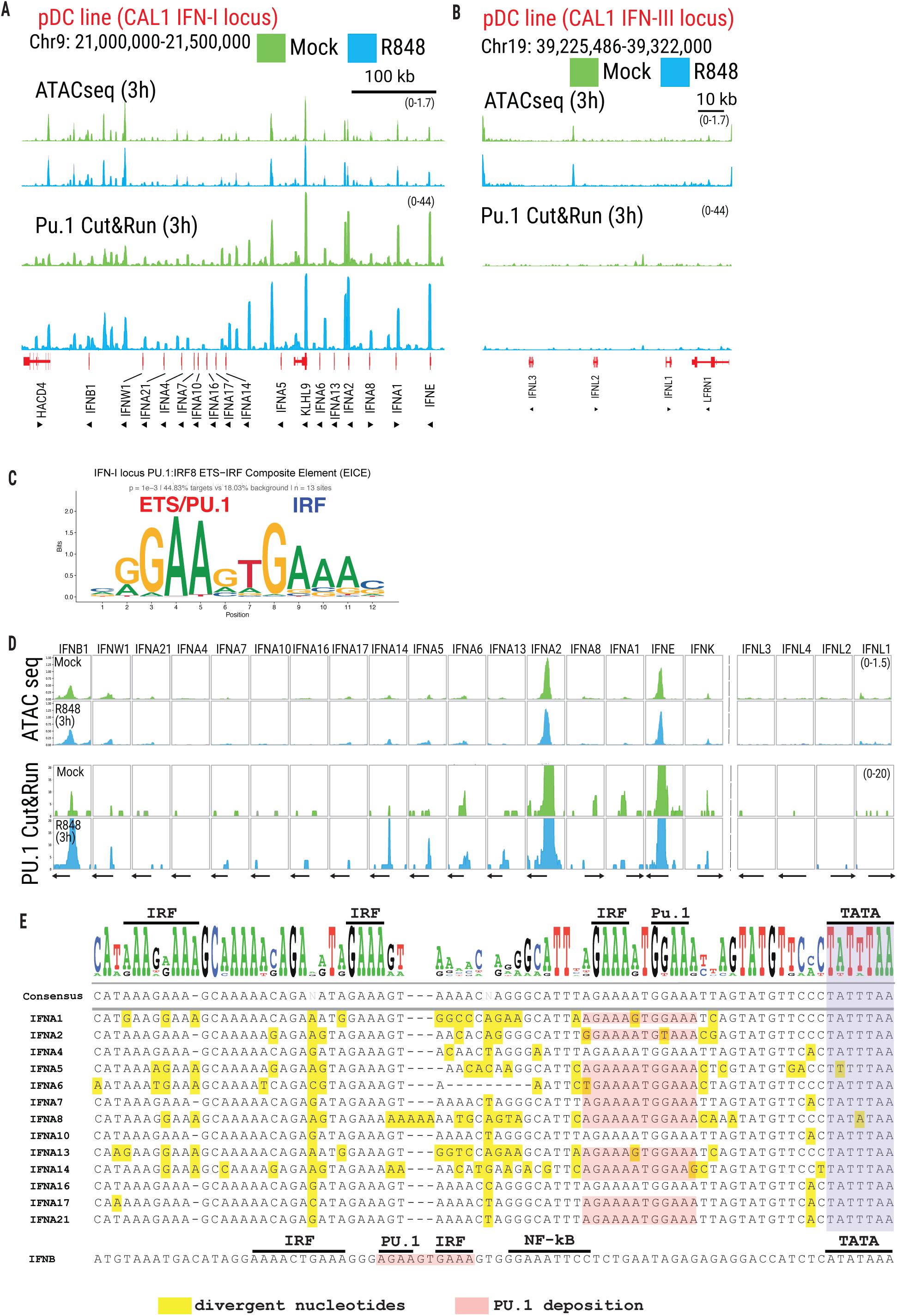
PU.1 occupancy overlaps with chromatin accessibility at the IFN-I locus. (A-B) Chromatin accessibility (top) and PU.1 CUT&RUN signal (bottom) in CAL-1 pDCs at the IFN-I locus (chr9: 21.0-21.5 Mb) **(A)** and IFN-III locus (chr19: 39,225,486-39,326,550) **(B)** at 3 h. **(C)** Sequence logo of canonical ETS-IRF composite elements (EICE) derived from HOMER enrichment analyses of ATAC-seq peaks at the IFN-I locus in CAL-1 pDCs (p = 1 ξ 10^-3^; 44.83% of target peaks vs 18.03% background; n = 19 sites). **(D)** Chromatin accessibility (top) and PU.1 CUT&RUN signal (bottom) at individual IFN-I (left) and IFN-III (right) gene promoters (1 kb upstream of TSS; gray dashed vertical lines indicate TSS position) in CAL-1 pDCs at 3 h. **(E)** Sequence alignment of IFN-I gene promoters showing conserved IRF, PU.1 (ETS), NF-ΚB and TATA-box elements. Yellow shading shows divergent nucleotides relative to consensus; pink shading indicates PU.1 CUT&RUN-supported occupancy sites. The IFNB1 promoter is shown separately for comparison. Mock (green) and R848-stimulated (blue). Data (A-B, D) are representative of one of 2 biological replicates.

### PU.1 binding sites are positioned in tandem with IRF elements at promoters within the IFN-I gene locus

Previous studies have defined the regulatory architectures of IFN-I promoters, revealing shared core elements as well as important distinctions (Supplementary Fig. 1A). The *IFNB1* promoter is organized into a well-characterized enhanceosome composed of four positive regulatory domains (PRD IV–III–I–II) ^52–57^, which are occupied by ATF/cJUN, IRF, and NF-κB family members and together mediate cooperative transcriptional activation. In contrast, *IFNA* gene promoters exhibit greater variability and tend to rely more heavily on IRF-driven regulation. *IFNW1* contains canonical ISREs and IRF motifs, whereas *IFNE* is regulated hormonally and through its 5′ untranslated region^58^. While examining PU.1 CUT&RUN deposition at the IFN-I promoter sequences, we identified a previously uncharacterized ETS family binding site within the *IFNB1* enhanceosome (Fig. 4C-E, Supplementary Table 2). This ETS motif lies between the two IRF-binding elements, IRFEi and IRFEii, in the proximal *IFNB1* promoter and precisely coincides with robust PU.1 occupancy. The close spatial arrangement of the ETS and IRF motifs defines an ETS-IRF composite element, a regulatory architecture previously described in immune genes such as *IGL*^59^, CYBB ^60^, *CD20*^61^, *NLRP3*^62^, and *IL18*^63^, but not previously reported at the *IFNB1* promoter. In addition to *IFNB1*, PU.1 deposition was observed at multiple *IFNA* gene promoters, with particularly strong binding at *IFNA2* and moderate enrichment at several additional *IFNA* genes (Fig. 4D-E). Promoter sequence analysis guided by PU.1 occupancy revealed that the IRF-binding site proximal to the TATA box of *IFNA* gene promoters also conforms to a PU.1-IRF composite element (Fig. 4D-E). These data suggested that PU.1 deposition is always in proximity to an IRF site in IFN-I promoters.

These observations led us to propose that PU.1 cooperatively binds IFN-I promoters together with an IRF family member in resting pDCs. Unlike IRF3, IRF5, and IRF7, which require activation-induced nuclear translocation, IRF8 is constitutively nuclear, highly expressed in pDCs, and known to physically interact with PU.1 through defined protein interaction domains. To test whether PU.1 and IRF8 bind IFN-I promoters in naïve pDCs, we performed electrophoretic mobility shift assays (EMSA) using nuclear extracts from mock-treated CAL-1 cells and radiolabeled promoter probes from *IFNB1*, *IFNL1*, and *IFNL2/3* (Fig. 5A). We observed strong binding of PU.1 and IRF8 to the *IFNB1* promoter probe, which was confirmed by a robust super shift upon addition of a PU.1 or IRF8-specific antibody (Figure 5B). In contrast, neither PU.1 nor IRF8 bound to IFNL promoter sequences, consistent with the absence of PU.1 deposition at the IFN-III locus (Fig 4B).

**Figure 5:**
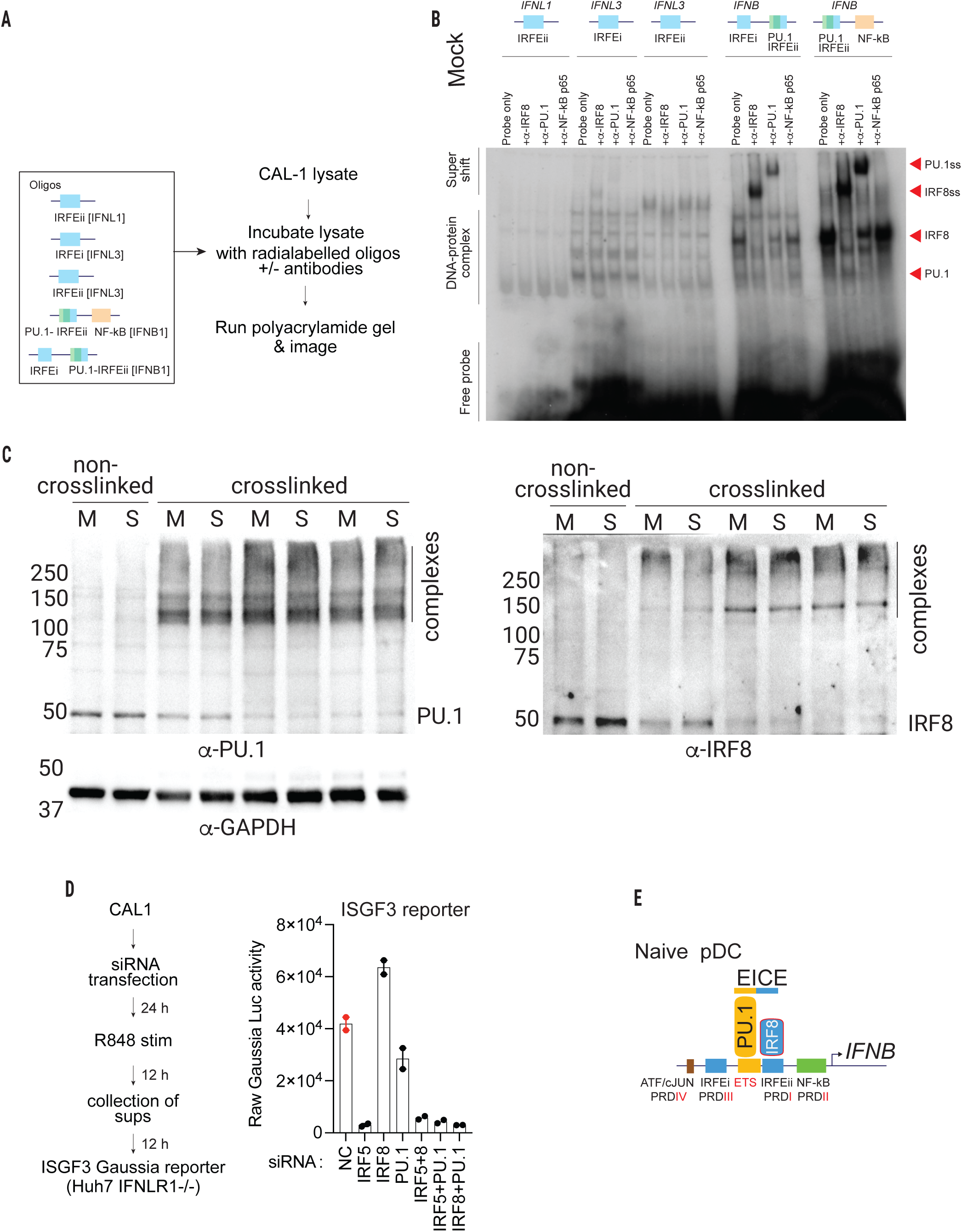
PU.1 and IRF8 interact at ETS-IRF-containing IFN gene promoters and are required for IFN-I induction in pDCs. **(A)** Schematic of the EMSA experiment design showing genomic sequences of radiolabeled oligonucleotide probes corresponding to IFNL1 IRFEii, IFNL3 IRFEi, IFNL3 IRFEii, IFNB1 (PU.1+NF-κB), and IFNB1 (IRFEi+PU.1+IRFEii) regulatory elements. IRF binding sequences are highlighted in blue, NF-ΚB binding sequences in orange, ETS/PU.1 binding sequences in green. **(B)** EMSA using cell lysate extracts from mock CAL1-pDCs. Antibodies against IRF8 (α-IRF8), PU.1 (α-PU.1), and NF-ΚB p65 (α-NF-ΚB p65) were used to identify DNA-bound protein complexes. Bands indicate supershift (PU.1ss, IRF8ss), DNA-protein complexes, and free probe. **(C)** Western blot analyses of PU.1 (left, α-PU.1) and IRF8 (right, α-IRF8) in non-cross-linked and DSS-cross-linked CAL-1 pDC cell lysate from mock (M) and R848-stimulated (S) conditions. **(D)** Gaussia luciferase activity in ISGF3 reporter cells (IFNLR1 knockout HUH7 cells) treated with supernatants from R848-stimulated CAL-1 pDCs nucleofected with siRNA targeting IRF5, IRF8, PU.1, IRF5+IRF8, IRF5+PU.1, and IRF8+PU.1 or negative control (NC) siRNA. The schematic shows the experiment design. **(E)** Schematic showing PU.1 and IRF8 co-occupancy at the EICE motifs within the IFNB1 enhanceosome in naïve pDCs, alongside ATF/cJUN (PRDIV), IRFEi (PRDIII), ETS, IRFEii, and NF-κB (PRDII) binding elements. Data in (E) are mean ± SEM, n = 3 biological replicates.

Together, these data identify previously unrecognized ETS-IRF composite elements within IFN-I promoters and demonstrate their occupancy by PU.1 and IRF8 in naïve pDCs.

To examine the physical interaction between IRF8 and PU.1 under naïve conditions, we performed endogenous co-immunoprecipitation (co-IP) assays using CAL-1 cell lysates. Cells were treated with DSS crosslinker to preserve protein-protein interactions, and the lysates were subjected to immunoprecipitation of endogenous PU.1 or IRF8. Immunoprecipitated complexes were resolved by SDS-PAGE and analyzed by immunoblotting for the presence or absence of the binding partner PU.1 or IRF8. We observed IRF8 and PU.1 co-immunoprecipitation under both basal and R848-stimulated conditions, indicating a physical interaction between these transcription factors in CAL-1 cells (Fig. 5C). This indicates a direct protein-protein interaction rather than simply binding to adjacent DNA elements. Previous studies have shown PU.1 and IRF8 interact through the IAD-PEST domain interactions^64^, consistent with the co-immunoprecipitation we observe in CAL-1.

To dissect the contribution of IRF5, IRF8, and PU.1 to type I interferon (IFN-I) gene regulation, we performed targeted knockdowns of IRF5, IRF8, PU.1, or their combinations in CAL-1 and assessed panIFN-I expression on an IFN reporter cell line (ISRE *Gaussia* reporter in an IFNLR1-/-background) in supernatants following R848 stimulation (Figure 5D). Silencing IRF5 abrogated panIFN-I production as it is one of the transcription factors required for IFN-I induction. In contrast, IRF8 knockdown increased panIFN-I expression relative to non-targeting controls, suggesting that IRF8 alone could act as a negative regulator of panIFN-I induction, presumably by competing with activating IRFs for the same recognition sequence. Combined knockdowns of IRF5 + IRF8, IRF5 + PU.1, or IRF8 + PU.1, led to a complete loss of panIFN-I induction (Figure 5D). These findings indicate that IRF5 is indispensable for IFN-I expression downstream of R848, whereas IRF8 and PU.1 together are critical for maintaining IRF5-mediated transcriptional activation. These data suggest that PU.1 and IRF8 complex occupy the ETS-IRF composite site within IFN-I promoters, maintaining the promoters in a transcriptionally poised state. This accessibility facilitates subsequent recruitment of activating IRFs (IRF3, IRF5, or IRF7) and NF-kB components for transcriptional activation of IFN-I. It is possible that activating IRFs could compete with IRF8, which is why we observe increased panIFN-I production during IRF8 knockdown.

These observations support a model in which lineage-restricted PU.1-IRF8 complexes (Fig. 5E) establish a poised chromatin that selectively licenses IFN-I, but not IFN-III, gene expression in pDCs.

### Intergenic ETS-IRF composite sites mark enhancer sites in the IFN-I gene locus in pDCs

Superimposition of ATAC-seq accessible sites and PU.1 deposition identified several overlapping peaks within intergenic regions of the IFN-I gene cluster, both at promoter and intergenic regions (Fig. 4A). We hypothesized that the PU.1 deposited accessible sites in the intergenic regions could serve as regulatory elements. Importantly, neither chromatin accessibility nor PU.1 deposition differed between mock and R848-stimulated conditions. To test if the PU.1-bound accessible regions possesses features of regulatory activity, we performed CUT&RUN for H3K4me1, H3K27ac, and RNA Pol II Ser5 phosphorylation proteins (PolIISer5p; Fig. 6A, Supplementary Table 2). Analysis of genomic regions spanning 5 kb upstream and downstream of IFN-I genes revealed at least thirteen sites where we observed deposition of PU.1 and enrichment for H3K4me1, H3K27ac, and Pol II Ser5p, all of which overlapped with ATAC-seq peaks (Fig. 6A). Together, the overlap of the above chromatin marks at PU.1-bound, accessible regions strongly suggests they are active enhancer elements. Deposition of H3K4me1 distinguishes enhancers from promoter regions, while enrichment of H3K27ac marks transcriptionally active enhancers. The Ser5-phosphorylated RNA Pol II deposition further supports enhancer function, as this deposition is associated with transcription initiation and enhancer RNA production at active regulatory elements. Additionally, motif analysis revealed that these candidate enhancer regions harbor ETS-IRF composite binding sites, suggesting that cooperative transcription factor binding potentially contributes to enhancer regulation at the IFN-I locus (Fig. 6B-D). Collectively, these findings indicate that the IFN-I gene cluster is embedded within a pre-configured enhancer landscape characterized by PU.1 occupancy and enhancer-associated chromatin features, thereby maintaining baseline chromatin accessibility and enabling rapid transcriptional responses upon stimulation. These data suggest that the some intergenic ETS-IRF composite elements potentially function as enhancer sites in IFN-I locus in pDCs. Enhancers loop around the promoter regions to increase the density of transcription factor complexes and co-activators to stimulate transcription. We propose that these enhancers elements are poised to enable rapid transcriptional responses upon stimulation. However, we did not observe this at the IFN-III locus, which remains inaccessible under both mock and R848 stimulation conditions.

**Figure 6:**
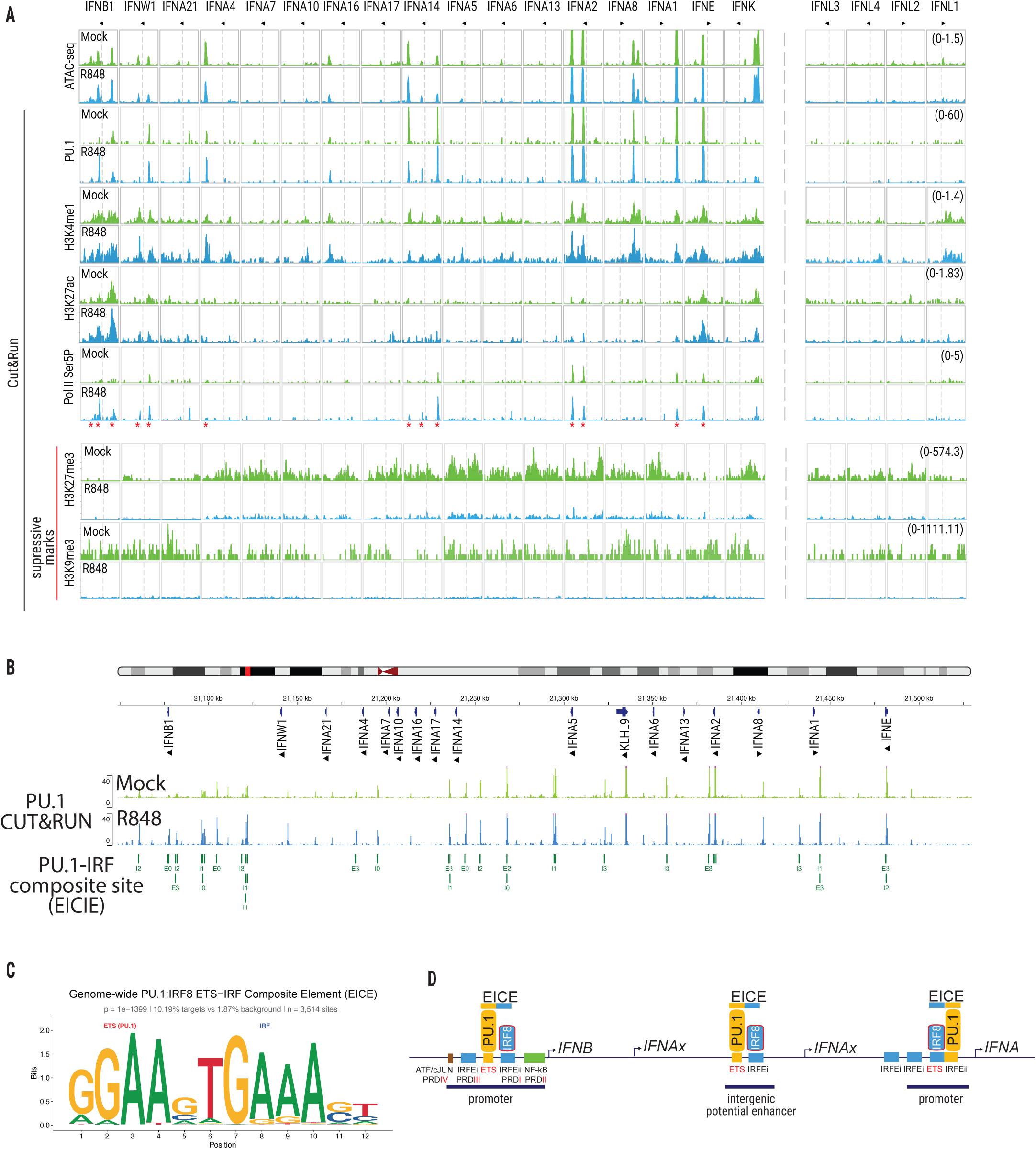
PU.1 constitutively occupies ETS-PU.1-containing promoters and enhancers across the IFN-I locus to establish a poised chromatin state for rapid IFN activation in pDCs. **(A)** Chromatin accessibility (ATAC-seq) and CUT&RUN signal of PU.1 H3K4me1, H3K27ac, Pol II Ser5P, H3K27me3, and H3K9me3 (at IFN-I and IFN-III loci in CAL-1 pDCs. Each track spans 5 kb upstream of the TSS through the gene body to 5 kb downstream; gray dashed vertical lines indicate TSS position. H3K27me3, and H3K9me3are shown as suppressive marks. Red asterisks show putative enhancer elements defined by co-occupancy of H3K4me1 and H3K27ac marks at intergenic ETS-IRF recognition sites. **(B)** IGV genome browser snapshot showing the distribution of ETS-IRF recognition sites across the IFN-I locus in CAL-1 pDCs. EICE sites are annotated by half-site orientation and spacer length: E = ETS half-site-centered (GGAA core); I = IRF half-site-centered (GAAA core); numeric suffix indicates spacer length in base pairs between half sites (0-3 bp). **(C)** Sequence logo of genome-wide EICE motif (p=1×10^⁻13⁹⁹^, 10.19% targets vs 1.87% background, n=3,514 sites). **(D)** Schematic of architecture at three classes of IFN gene regulatory elements: IFNB1 promoter (left), IFNAx intergenic potential enhancers (center), and IFNAx promoters (right). Each element contains ETS-IRF recognition sites occupied by both PU.1 and IRF transcription factors. Mock (green) and R848-stimulated (blue, 3 h). Data in (A) are representative of one of the 2 biological replicates.

### Delayed chromatin accessibility at the IFN-I locus in epithelial cells

Epithelial organoid cells lacked chromatin accessibility peaks at the IFN-I locus at 6 h, in both resting and RIG-I ligand-stimulated conditions. In contrast, the IFN-III locus was accessible across all time points in both mock and stimulated cells (Fig. 1). A similar expression pattern was observed in a hepatocyte cell line that tolerated extended periods of PAMP stimulation up to 24 h. Epigenetic silencing of the IFN-I locus in epithelial cells was further supported by the presence of H3K9me3 and H3K27me3 repressive marks (Supplementary Fig. 2).

However, at 12 and 24 h post-stimulation, moderate chromatin accessibility was observed at *IFNB1, IFNW1, IFNA4, IFNA7,* and *IFNA10* in stimulated conditions, mirroring IFN-I gene expression kinetics in these cells. Interestingly, pDC-associated IFN genes, *IFNA2, IFNE, IFNA14, IFNA5*, and *IFNA6*, remained inaccessible at all time points regardless of stimulation. The IFN-III locus was accessible in both mock and stimulated conditions at all time points examined. Despite this constitutive accessibility, IFN-III expression was only detected after 6 h of stimulation, implicating promoter strength as a rate-limiting determinant of transcriptional initiation, as shown in Fig. 3H. Taken together, these data indicate that the delayed IFN-III gene expression observed despite an open chromatin configuration is attributable, at least in part, to intrinsically weaker promoter activity driving IFN-III gene expression.

## Discussion

The promoters of IFN-I and IFN-III genes have been extensively characterized. The *IFNB1* promoter operates through a well-defined PRD IV–III–I–II architecture occupied cooperatively by ATF/c-JUN, IRF, and NF-κB family members, while *IFNA* gene promoters rely predominantly on tandem IRF modules (IRFi–IRFii–IRFiii) for activation^52–57^. The activating transcription factors that bind these elements, principally IRF3, IRF5, IRF7, and NF-κB, have been identified across many cell types. Yet a fundamental question that remains unresolved is why pDCs produce copious IFN-I within hours of TLR engagement, whereas other nucleated cells capable of TLR, RLR, and DNA sensing, including epithelial cells, mount a delayed and comparatively modest IFN-I response. Rapid induction is currently explained by constitutively high basal expression of IRF5 and IRF7 in pDCs, which enables immediate transcriptional activation without the IRF7 induction step required in most other cell types. While IRF5 and IRF7 dosage are important, they do not fully account for differences IFN subtype production between cell types. Epigenetic regulation could explain these differences, yet studies of chromatin regulation in interferon biology have focused almost exclusively on the *IFNB1* gene^65–68^, and a locus-wide cell-type-specific epigenetic analysis of human IFN gene clusters has not been undertaken to date. In this study, we show that the IFN-I and IFN-III loci carry opposing, constitutively established chromatin accessibility landscapes in pDCs and epithelial cells, and that this locus-wide epigenetic divergence is enforced by lineage-specific ETS-IRF regulatory architecture and repressive histone marks, primary determinants of cell-type-specific IFN gene induction (Fig 7).

**Figure 7.**
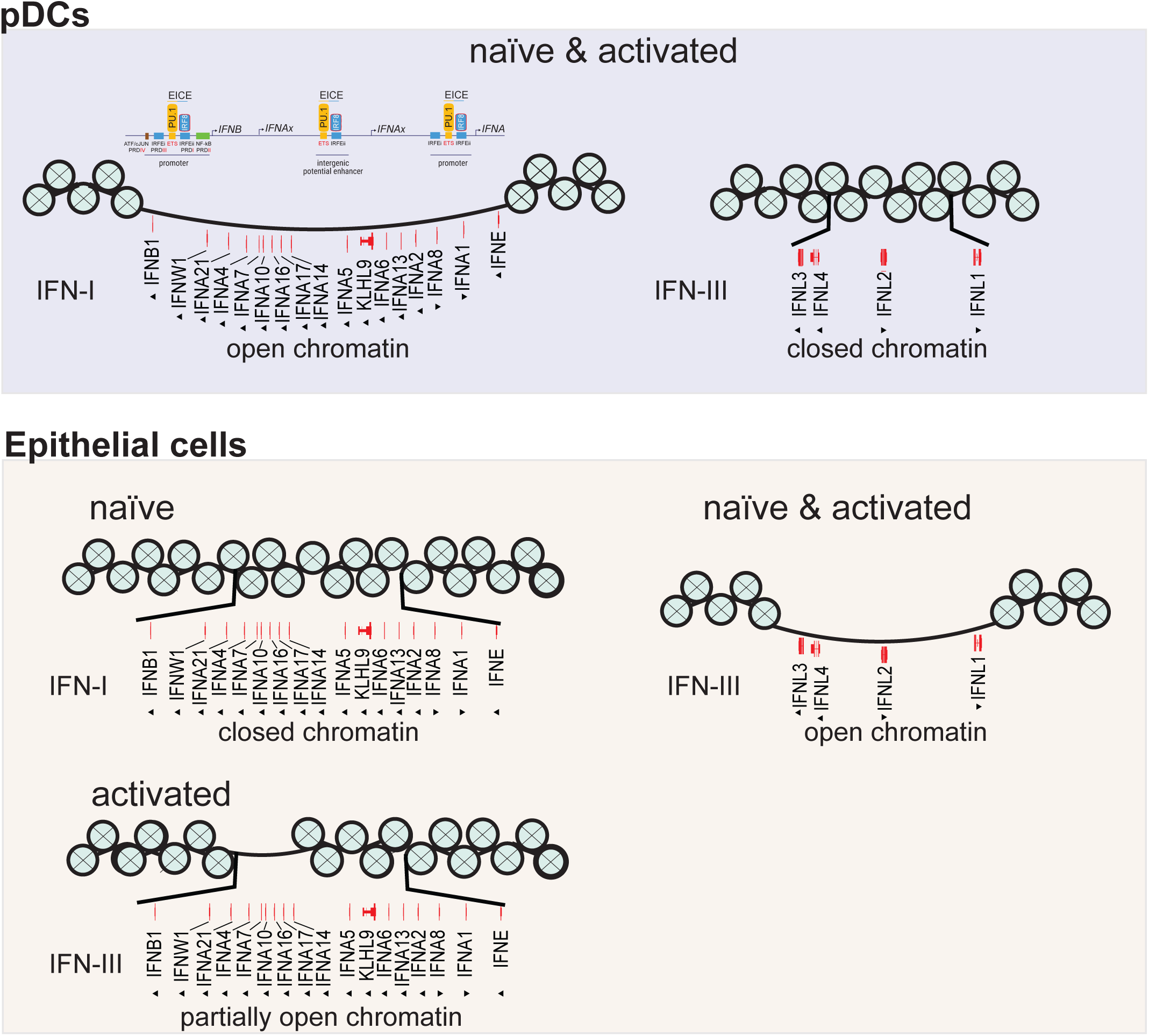
Chromatin accessibility and epigenetic landscape governing IFN-I and IFN-III induction in pDCs and epithelial cells. In plasmacytoid dendritic cells (pDCs), PU.1 and IRF8 constitutively co-occupy EICE motifs across the IFN-I locus at both promoters and intergenic putative enhancers in naïve and activated states, maintaining an open chromatin configuration that poises the locus for rapid transcriptional activation upon PAMP stimulation. The IFN-III locus, which lacks PU.1 occupancy in pDCs, remains in a closed chromatin state. In intestinal epithelial cells, the IFN-I locus adopts a closed chromatin configuration, while the IFN-III locus is selectively accessible and inducible. Together, these data support a model in which lineage-specific transcription factors, particularly PU.1 in pDCs, determine locus-specific chromatin accessibility and epigenetic landscape, thereby specifying whether a cell preferentially produces type I or type III interferons in response to innate immune stimulation. Circles represent nucleosomes; red marks indicate ETS-IRF recognition sites.

In pDCs, we show that the IFN-I locus is constitutively open and transcriptionally poised, in stark contrast to the IFN-III locus, a distinction determined by PU.1-IRF8 binding. Analysis of IFN-I proximal promoters revealed ETS-IRF composite elements at the PRDIV site of the *IFNB1* promoter and IRF motif proximal to the TATA box in *IFNA* genes. Consistent with these motif arrangements, we show that PU.1 and IRF8 bind to IFN-I promoters but not to IFN-III promoters, which lack ETS (PU.1) recognition sequences. These data provide a mechanistic basis for selective locus poising, in which the ETS-IRF composite element is required for PU.1-IRF8 occupancy, and the absence of this element from the IFN-III locus explains the lack of accessibility in pDCs.

The poising function of ETS-IRF composite motifs appears to extend beyond individual promoters. ETS-IRF composite motifs are distributed across the intergenic regions of the IFN-I locus, and we identify candidate enhancer elements at these sites based on co-occupancy of PU.1, active histone marks (H3K4me1, H3K27ac), and RNA Pol II. We propose that this network of ETS-IRF composite-element-anchored enhancers maintains the locus in an open, primed configuration, enabling the rapid, robust IFN-I response that is the hallmark of pDC biology. This enhancer-promoter interactions may further depend on the three-dimensional organization of the locus, where cohesin-dependent chromatin looping, which has recently been shown to reorganize the IFN-I locus during pDC activation in murine pDCs^69^, and it is plausible that PU.1-IRF8-occupied ETS-IRF composite enhancers serve as loop anchors that bring specific regulatory elements into contact with their target gene promoters in a stimulation-dependent manner. While previous studies have identified the transcription factors that activate IFN gene transcription, our work defines how IFN-I promoters and the broader locus are maintained in a poised state by PU.1-IRF8 before the activating transcription factors are engaged. Further analysis of enhancer-enforced IFN-I subtype expression is warranted to understand how PU.1 cooperates with IRF8 in certain stimulatory in pDCs. Dysregulation of PU.1-IRF8 may contribute to ineffective viral control or autoimmunity, as genetic polymorphisms in IRF8 are associated with Systemic Lupus Erythematosus^70,71^.

In epithelial cells, the IFN-I locus displays a fundamentally distinct organization. Constitutive repression, enforced by H3K9me3 and H3K27me3 marks, keeps the locus inaccessible at rest and during early stimulation, with accessibility emerging selectively at 12-24 h and only at a restricted subset of IFN-I genes (*IFNB1*, *IFNW1*, *IFNA4*, *IFNA7*, *IFNA10*). The entire IFN-I locus and the pDC-associated *IFNA* subtypes, which rely most heavily on the ETS-IRF composite architecture for poising, remain silenced in epithelial cells, a finding consistent with the absence of PU.1 and IRF8 expression outside the hematopoietic lineage. The late and partial chromatin accessibility we detect at a subset of IFN-I genes, *IFNB1*, *IFNA4*, *IFNA7*, and *IFNA10*, at 12-24 h post-stimulation is consistent with a previous study showing stimulus-driven removal of repressive H3K9me2, deposited by G9a (EHMT2), at *IFNB1* in non-immune cells^72^. We propose the permanently inaccessible pDC-associated *IFNA* subtypes likely carry a deeper repressive architecture enforced by H3K9me3 deposition, which is refractory to stimulus-driven remodeling in the epithelial lineage. The delayed accessibility observed at the IFN-I locus in epithelial cells may also partly reflect a dependency on stimulus-driven chromatin remodeling, wherein IRF3 and NF-ΚB must first assemble an enhanceosome that recruits SWI/SNF remodeling complexes to displace the positioned nucleosome at the *IFNB1* transcriptional start site, a prerequisite step that itself imposes a delay before productive RNA Pol II engagement (reviewed in ^73^).

Together, these observations reframe the cell-type specificity of IFN-I production not as a quantitative difference in signaling capacity but as a qualitative difference in locus architecture established prior to any stimulus^73^.

The IFN-III locus is constitutively open in epithelial cells across all conditions examined. It supports only delayed transcriptional induction, partly due to its promoter strength, as we show substantially weaker intrinsic activity at *IFNL* promoters than at *IFNB1* following SeV infection. Chromatin accessibility is necessary but not sufficient for an early interferon (IFN) response. The strength of the core promoter imposes an additional, independent transcriptional constraint. These data suggest two independent mechanisms regulating IFN gene expression.

Chromatin accessibility determines whether a gene can be activated, and intrinsic promoter strength, which governs the rate of gene expression.

However, the IFN-III locus is not accessible in either naive or activated blood-derived pDCs. We propose that this is due to the absence of ETS-IRF composite sites on the promoter. It is worth noting that the closed IFN-III locus we observe in pDCs may not be absolute across all tissue contexts. Since TGF-β induces transcription of *IFNL* genes in murine gut-resident pDCs, an effect not observed in splenic pDCs, this suggests that local tissue signals can override lineage-encoded locus silencing in a compartment-specific manner^74^. This raises a possibility that the chromatin architecture of the IFN-III locus in pDCs may not be fixed but may be subject to tissue specific reprogramming^74^.

A current limitation of our study is the absence of sensitive antibodies compatible with CUT&RUN for IRF3, IRF5, IRF7, and IRF8, which prevents us from determining the temporal dynamics of transcription factor(s) exchange at the IFN-I locus following pDC activation. Specifically, it remains unclear whether PU.1 and IRF8 are ejected upon activation to make way for incoming IRF3, IRF5, or IRF7, or whether they are retained and incorporated into the activating transcriptional complex. The latter possibility is supported by prior evidence showing that IRF5 and IRF8 are in physical proximity following CpG stimulation in pDCs^46^, suggesting that the PU.1-IRF8 complex may persist at IFN-I promoters in the activated state rather than being displaced. Broadly, resolving the dynamics of activating and poised transcription factor occupation will require the development of high-sensitivity antibodies or epitope-tagged knock-in cell models suitable for single cell chromatin profiling and represents an important direction for future work.

Collectively, our findings support a model in which distinct IFN programs of pDCs and epithelial cells are influenced by chromatin architecture and *cis*-regulatory sequences, lineage-specific TF binding, repressive and active histone modifications, and intrinsic promoter strength. In combination, these factors determine how a cell responds to activating signals, clarifying how cell-type identity constrains innate immune responses independently of signaling pathway activation. An important open question is whether the candidate enhancer elements we identified at intergenic ETS-IRF composite sites in pDCs regulate the expression of specific *IFNA* subtypes. Our data, on pDCs or epithelial cells, capture a snapshot in a broader functional context of these cells subjected to distinct tissue-specific cues and pathogens, potentially licensing distinct IFN-I or IFN-III subtype programs tailored to the local tissue or immune context. Future work using enhancer perturbation and single-cell approaches will be needed to resolve this subtype specificity and its functional consequences for antiviral restriction and autoimmunity.

## Methods

Reagent specifics are presented in Supplementary Table 3.

### Primary cells and cell line cultures

Human blood was obtained from anonymized healthy donors through Bloodworks Northwest. Peripheral blood mononuclear cells (PBMCs) were isolated within 4h of phlebotomy. PBMCs were isolated from EDTA-anticoagulated whole blood by density gradient centrifugation using Lymphoprep (Stem Cell technologies, #18060) at 800 g without a brake. Cells were further purified by negative selection using the EasySep Direct Human PBMC Isolation Kit (Stem Cell Technologies, #19654) according to the manufacturer’s instructions.

Fresh PBMCs were used to isolate pDCs using the EasySep Human Plasmacytoid DC Isolation Kit (Stem Cell technologies, #17977). Primary pDCs and CAL-1 cells (pDC cell line) were cultured in RPMI 1640 (Invitrogen) supplemented with 10% fetal bovine serum. PH5CH8, HuH7 reporter, and HEK293 cells were cultured in DMEM with 10% fetal bovine serum, 1% HEPES, 1% nonessential amino acids (NEAA), 1% sodium pyruvate, and 1% penicillin-streptomycin-L-glutamine. All cells were cultured at 37 °C temperature in a humidified incubator with 5% CO₂.

### Human enteroid culture

Human duodenal enteroids were derived from anonymized healthy donor tissue from three donors (HD003 - 68-year-old male, HD005 - 40-year-old male, HD007 - 70-year-old female) obtained from surgical resection, as previously described for human ileal enteroids^75^. For continuous culture, enteroids were maintained in Matrigel domes in WRNE medium consisting of Advanced DMEM/F12, N-acetyl-L-cysteine (1 mM, Sigma), HEPES (10 mM, VWR), GlutaMAX (1x), B-27 (1x) supplement, NEAA (1x), 1% antibiotic-antimycotic (Sigma), 50% L-WRN conditioned medium (CM), mouse epidermal growth factor (mEGF, 50 ng ml⁻¹), SB202190 (10 µM, Sigma), human gastrin (10 nM, Sigma), nicotinamide (10 mM), and A 83-01 (500 nM, Tocris). In addition, cells were supplemented with PGE2 (100 nM, Cayman) at the time of passaging, and Y-27632 (10 µM, Hello Bio) was added to the medium when cells were first plated after passaging. L-WRN cells^76^ were obtained from Thaddeus Stappenbeck (Cleveland Clinic) and propagated in DMEM with 10% FBS, GlutaMAX (1x), G418 (0.5 mg mL^-1^), and hygromycin (0.5 mg ml⁻¹). Once cells were >95% confluent, medium was changed to 20% FBS and GlutaMAX (1x) in Advanced DMEM/F12. Cells were incubated for 6 days with daily medium collection and replacement. The resulting CM from all days was pooled and filtered. In addition, cells were supplemented with PGE2 (100 nM, Cayman) at the time of passaging, and Y-27632 (10 µM, Hello Bio) was added to the medium when cells were first plated after passaging.

To generate monolayers of cells, human duodenal enteroids were removed from Matrigel using cell recovery solution; dissociated in 0.05% trypsin/0.5 mM EDTA at 37°C for 6-10 min; quenched with DMEM containing 10% FBS, 10 mM HEPES, 1% NEAA, GlutaMAX (1x), and Y-27632 (10 µM); and mechanically dissociated with a pipette. Cells were filtered through a 40 µm cell strainer, centrifuged at 400 g for 5 min, and resuspended in WRNE medium containing Y-27632 (10 µM) but without nicotinamide. 2.0-2.8 x 10⁵ cells/well, depending on the enteroid line, were plated onto 96-well plates previously coated with human placental collagen, type IV. Cells were switched to WRNE lacking both Y-27632 and nicotinamide one day after seeding. The cultures were incubated for five days prior to analysis, with medium changes every two days. After 5 days, cells were rested overnight with 0.1% FBS and GlutaMAX (1x) in Advanced DMEM/F12. HCV-PAMP (500 ng mL^-1^) was transfected using Mirus TransIT-X2 to stimulate cells for 6 h after replacing the media with regular 20% FBS and GlutaMAX (1x) in Advanced DMEM/F12.

### Interferon expression

pDCs and CAL-1 cells were stimulated with R848 (1 µg ml⁻¹), and PH5CH8 and intestinal epithelial cells derived from human enteroids were stimulated with HCV-PAMP (500 ng ml⁻¹). HCV PAMP was complexed with Mirus TransIT-X2 at a 1:3 ratio (w/v) in Opti-MEM culture medium for 20 min at room temperature, and a total of 500 nanograms of HCV PAMP was used for each 10^6^ cells. Supernatants were harvested at different time points (0, 3, 6, 12, 24, 36, and 48 hours) after stimulation. For interferon measurement, 50 µl of supernatants were transferred to reporter cells lacking IFNAR1 or IFNLR1 receptors and incubated for 24 h. Gaussia luciferase-based promoter activity was measured using Gaussia glow assay buffer and substrate according to the manufacturer’s instructions. Data included shows mean ± SEM. GraphPad Prism version 10.6.1 was used for statistical analyses. The differences in IFN signaling kinetics between Ph5Ch8 and Cal-1 cells were analyzed using ordinary two-way ANOVA, with time and cell type as the two components, including the interaction term. Post-hoc comparisons were done using Šídák’s multiple comparisons test to compare Ph5Ch8 and Cal-1 at each individual timepoint (simple effects within rows). To compare different timepoints’ kinetics, Dunnett’s multiple comparisons test was used within each cell type separately (simple effects within columns), using 24 h as the reference timepoint for PH5CH8 and 3 h as the reference timepoint for Cal-1, corresponding to their respective expression peaks. A p-value of <0.05 was considered statistically significant.

### Luciferase promoter construct design and cloning

The proximal promoter regions of *IFNB1*, *IFNL1*, and *IFNL2/3* were synthesized incorporating flanking restriction sites. Amplified fragments were cloned into the multiple cloning site of the pGL3.1 firefly luciferase reporter vector (Promega) in the sense orientation. IRF binding site mutations were designed to disrupt the core GAAA IRF recognition sequence at each predicted IRF element within the *IFNB1* PRD-I and PRD - III, *IFNL1*, and *IFNL2/3* promoters. NF-κB binding site mutations were introduced at the PRDII element of the *IFNB1* promoter and at the corresponding NF-κB sites in *IFNL* promoters by substitution of the core κB consensus sequence. All constructs were verified by sequencing. A promoter-less pGL3.1 vector (Basic) was included in all experiments as a negative control to determine basal luciferase background activity. Mutant and wild-type constructs were tested in parallel in all experiments.

### Cell transfection and viral infection

HEK293T cells or the relevant cell line were seeded at 2 × 10⁵ cells per well in 24-well plates 24 h prior to transfection. Cells were transfected using transfection reagent, Mirus TransIT-X2, according to the manufacturer’s protocol, using 100 ng of firefly luciferase reporter plasmid and 10 ng of a pMaxGFP expression plasmid per well. The pMaxGFP plasmid was co-transfected as an internal transfection efficiency control. Total plasmid DNA was kept constant across conditions by supplementing with empty vector where required. At 24 h post-transfection, cells were either left as mock-infected controls or infected with Sendai virus (SeV; Cantell strain) at a multiplicity of 60 hemagglutination units per milliliter (60 HAU/mL). Cells were harvested at 24 h post-infection for luciferase and GFP quantification.

### Luciferase and GFP quantification

Cells were lysed in passive lysis buffer (Promega) for 15 min at room temperature with gentle rocking. Firefly luciferase activity was measured using the Luciferase Assay System (Promega) on a luminometer according to the manufacturer’s instructions. GFP fluorescence was measured in the same lysates using a fluorescence plate reader (excitation 488 nm, emission 507 nm) to quantify transfection efficiency. Raw firefly luciferase values were normalized to GFP fluorescence intensity for each well to correct for differences in transfection efficiency. Normalized luciferase values are reported as relative light units (RLU) normalized to GFP (RLU/GFP). All assays were performed in biological triplicate with technical duplicates, and data are presented as mean ± SEM.

### siRNA-based knockdown and stimulation

CAL-1 cells were nucleofected with siRNA targeting IRF5 (Thermo Fischer Scientific, s7514), IRF8 (Origene, SR302298), and PU.1 (Santa Cruz, sc36330), individually or in combination, along with a non-targeting siRNA control. 1 x 10^6^ cells were suspended in SF cell line solution supplemented with 3.5 μl of supplement (Lonza) and siRNA at a final concentration of 20-50 nM. Nucleofection was performed on Lonza 4D Nucleofector (SF kit) with program DN100. Cells were immediately transferred to a pre-warmed complete RPMI medium and allowed to recover for 24 hours. Cells were then stimulated with R848 (1 µg ml⁻¹) for 24 h prior to downstream assays.

### Plasmid transfection and overexpression studies

HEK293 cells were seeded the day before transfection to 50-70% percent confluence. Cells were transfected with expression plasmids or empty vector controls using Mirius Trans-IT X2 transfection reagent as per the manufacturer’s instructions using a total of 100 ng of plasmid DNA in a 96 well format. Cells were harvested at 36 h post-transfection for RNA and protein analyses. Overexpression was verified by western blot.

### Western blot

Cells were harvested and washed once with PBS and lysed with 50 µL of RIPA buffer (10 mM Tris pH 7.5, 150 mM NaCl, 1% Triton X-100, 0.5% NaDOC) supplemented with Halt Protease and Phosphatase inhibitor cocktail (1%). Protein was clarified by centrifugation at 10,000 g for 10 min at 4 °C, and concentration was determined by a Bradford assay (Bio-Rad). Equal amounts of protein lysate were loaded in 1× Laemmli buffer with 2.5% β-mercaptoethanol (BME) and boiled at 95 °C for 5 min. Protein samples were separated by SDS-PAGE in Tris-glycine gels (Bio-Rad) and transferred to a nitrocellulose membrane (GVS Filter Technology) in Towbin buffer (25 mM Tris, 192 mM glycine, 20% methanol, 0.025% SDS). Transferred membranes were then blocked in 3% BSA in tris-buffered saline with 0.1% Tween-20 (TBST). Membranes were incubated with primary antibodies in 3% BSA in TBST overnight at 4 °C, washed three times for 3 min in TBST, and incubated with species-specific secondary antibodies conjugated to horseradish peroxidase for 1 h. Membranes were washed again and incubated with ECL substrate, then chemiluminescence was detected using a Bio-Rad ChemiDoc XRS+ imaging instrument.

### Electrophoretic mobility shift assay (EMSA)

EMSA was carried out as previously described^39^. Briefly, nuclear extracts were prepared from CAL-1 cells using the CellLytic NuCLEAR extraction kit (Sigma-Aldrich). Protein concentration was measured with a Bio-Rad protein assay and samples were stored at –70°C until use. DNA oligonucleotides corresponding to the predicted IRF/PU.1/NF-kB binding sequences of the *IFNL1/3 (same sequence)*, and *IFNB1* genes were synthesized (Integrated DNA Technologies). Annealed double-stranded probes were labeled with [α-^32^P] deoxycytidine triphosphate (3000 Ci/mmol; PerkinElmer Life and Analytical Sciences) by fill-in using the Klenow fragment of DNA polymerase I (Invitrogen). ^32^P-labeled double-stranded oligonucleotides were purified using mini–Quick Spin Oligo Columns (Roche Diagnostics). DNA-protein binding reactions were performed in a 20-µL mixture containing 5 µg nuclear protein and 1 µg poly(dI-dC) (Sigma-Aldrich) in 4% glycerol, 1 mM MgCl_2_, 0.5 mM ethylenediaminetetraacetic acid, 0.5 mM dithiothreitol, 50 mM NaCl, 10 mM Tris-HCl (pH 7.5). Nuclear extracts were incubated with 1 µL ^32^P-labeled oligonucleotide probe (10,000 cpm) at room temperature for 20 minutes and then loaded on a 5% polyacrylamide gel (37:5:1). Electrophoresis was performed in 0.5x TBE for 2 hours at 130 V, and the gel was visualized by autoradiography. For antibody supershift experiments, nuclear extracts were incubated with 2 μL of antibody for 1 h on ice before the addition of ^32^P- labeled DNA probe.

After the addition of labeled DNA probe, the binding reaction was incubated for an additional 20 min at room temperature. The antibodies used were anti-IRF8, anti-PU.1, and anti-NFkB (Santa Cruz Biotechnology Inc.).

### ATAC-seq

Chromatin accessibility was assessed by ATAC-seq using the Active Motif ATAC-Seq Kit (Active Motif, cat. No. 53150), according to the manufacturer’s instructions and established protocols^42^. 5 × 10⁴ cells (pDCs, CAL-1, human intestinal organoids, and PH5CH8) were harvested under mock and stimulated conditions, nuclei were isolated, and transposition was performed using Tn5 transposase at room temperature for 30 min. DNA was purified and PCR-amplified to create sequencing libraries; library quality and fragment size distribution were assessed on Agilent 4200 TapeStation before conducting paired-end sequencing on the Illumina NovaSeqX Plus platform.

ATAC-seq reads were processed using the nf-core/atacseq pipeline (v2.1.3; containerized in Docker). Adapter sequences were removed with TrimGalore/Cutadapt, and reads were aligned to the human reference genome (GRhg38/hg38 primary assembly) with Bowtie2 (v2.3.5^77^) with default parameters. Mitochondrial reads were removed, and PCR duplicates were excluded using Picard MarkDuplicates. Peaks were called per replicate using MACS2 (V2.1.1; Zhang *et al*., 2008) with paired-end setting (-f BAMPE) at q<0.01. Removal of peaks overlapping ENCODE hg38 blacklist regions was done to remove artifactual high signals^77^ using default parameters. Mitochondrial reads were removed and PCR duplicates were excluded using Picard MarkDuplicates. Tn5 insertion sites were offset by +4/−5 base pairs, and genome-normalized bigwig files were created using deepTools bamCoverage^78^. Removal of peaks overlapping ENCODE hg38 blacklist regions was done to remove artifactual high signals^79^. Quality metrics were calculated using MultiQC^80^. Genome-normalized bigwig files were generated using deepTools banCoverage. bedGraph files were used for locus-level visualization in SparK(v2.0). pDC ATAC-seq was performed on cells from two distinct healthy donors.

ATAC-seq studies for CAL-1 and PH5CH8 were performed in two independent biological replicates. Intestinal organoid-derived IEC ATAC-seq was performed from two donors; data from one donor were discarded after quality control assessment, and data from one donor are shown. Representative tracks of ATAC-seq signal and peak distributions are shown.

### Cut and Run

For transcription factors and epigenetic marks, CUT&RUN^81^ was performed in CAL-1 and PH5CH8 cells under mock and treated conditions. Briefly, cells were immobilized on concanavalin A-coated magnetic beads, permeabilized with digitonin, and incubated with anti-SPI1/PU.1 antibody overnight at 4°C prior to AutoCUT&RUN processing. Libraries were prepared using a Beckman Biomek i7 liquid handling platform. End-repair, adapter ligation, and PCR amplification reactions were performed, and purified PCR products were analyzed on an Agilent 4200 TapeStation. Libraries were quantified and pooled at equimolar concentrations for paired-end 50 bp sequencing on the Illumina NovaSeq X Plus platform. Antibodies used for CUT&RUN included anti-SPI1/PU.1, anti-IRF8, anti-H3K27ac, anti-H3K4me1, anti-H3K27me3, and anti-RNA Pol II Ser5P/2P (Supplementary Table 3).

Base calls were converted to FASTQ files on Illumina BaseSpace. Adapter trimming was performed using Trim Galore/cutadapt. Reads were aligned to the human reference genome (GRCh38/hg38, primary assembly) using Bowtie2^77^ or the R Subread aligner, depending on the experiment. Mitochondrial reads were removed, PCR duplicates were marked with Picard (v1.134), and secondary and supplementary alignments were excluded. Quality-control metrics for sequencing, alignment, and peak-calling steps were aggregated using MultiQC. For peak calls, macs3 v3.0.1 was run in paired-end mode with a q-value cutoff of 0.05 on filtered reads and additionally with SEACR v1.3 in stringent mode with an empirical FDR threshold of 0.01 against matched IgG controls^82^. pDC and CAL-1 CUT&RUN experiments were performed in two independent biological replicates; one representative replicate is shown per condition. Signal tracks represent spike-in (*E. coli* K12) normalized read depth. IgG controls were used for SEACR peak calling background subtraction.“

Transcription factor motif enrichment was assessed using HOMER (findMotifsGenome.pl) with 200 bp windows centered on peak summits and automatically generated GC-content and length-matched background sequences obtained from hg38^83^. Known motif enrichment was obtained from the HOMER built-in motif database using Benjamini-Hochberg multiple testing correction; enriched motifs were considered enriched at q ≤ 0.01. The motif enrichment analyses were performed genome-wide and restricted to IFN-I (chr9:21,000,000 - 21,500,000) and IFN-III (chr19:39,225,486 - 39,326,550) loci of primary pDCs, CAL-1, PH5CH8 and IECs.

Please note that we restricted the loci to a smaller genome fragment, and for the loci that yielded insufficient accessible peaks, ATAC signals were extracted from tabix-indexed bedGraph files restricted to the locus, and the accessible region was defined using MACS3 bdgpeakcall^84^ with a minimum length of 100 bp, maximum gap of 200 bp, and signal cutoff thresholds of 5-15 adjusted per condition^84^. For pDC and CAL1, ATAC-seq at the IFN-I locus, peaks were derived directly from genome-wide MACS2 narrowPeak calls which were filtered to the IFN-I and IFN-III locus coordinates; results from fewer than 30 peaks are interpreted with appropriate caution. Motif enrichment analysis was not performed at the IFN-III locus in pDCs and CAL-1, or at either locus in PH5CH8 and IECs, due to insufficient peak numbers (fewer than 20 peaks) for reliable enrichment estimation.

Statistical analyses and data visualization were performed in R (v4.5.1) and Python (v3.13.2) using the tidyverse package collection (v2.0.0^85^) and numpy/pandas libraries^86,87^, respectively. Sequence logos were generated using ggseqlogo (v0.2.2^88^) with y-axis scaled to information content (bits).

Genome-wide chromatin and epigenetic signal tracks and peak distributions were plotted using SparK (Kurtenbach, S. & Harbour, J.W. SparK (v2.0), pyGenomeTracks^89^, and Integrative Genome Viewer (IGV^90^). For analyzing and plotting genomic data R (4.5.1), Python (3.13.2), tidyverse (2.0.0), ggplot2 (4.0.1), ggseqlogo (0.2.2)^88^, DiffBind (3.18.0), GenomicRanges (1.60.0^91^), ChIPseeker (1.44.0^92^), clusterProfiler (4.16.0^93^) were used. All ATAC seq and Cut and Run experiments were done in duplicates, and signals were normalized over IgG background; one representative replicate is presented in the Figures. GraphPad Prism was used for bar graphs and kinetics plots. All command-line parameters, software versions, and configuration files for nf-core pipelines can be requested from the corresponding author.

## Data and code deposition

All sequencing data (ATAC-seq and CUT&RUN) created in this study have been deposited in the NCBI Gene Expression Omnibus (GEO:GSE334179).

## Declaration of generative AI and AI-assisted technologies in the writing process

Claude AI was used in the writing process only to improve the readability and editing language of the manuscript. After using Claude AI, we have reviewed and edited the content as needed and take full responsibility for the content of the publication.

## Supplementary Figure and Table Legends

**Figure S1: (A)** Schematic of IFN promoters. **(B)**Schematic and genomics sequences of regulatory elements of IFNL1 IRFEii, IFNL3 IRFEi, IFNL3 IRFEii, IFNB1 (IRFEii+NF-κB), and IFNB1 (IRFEi+IRFEii). IRF binding sequences are shown in blue, NF-ΚB binding sequences in orange and ETS/PU.1 binding sequences in green. EMSA probes used experiments in Figure 5B are based on these promoter sequences.

**Figure S2: Epigenetic landscape at IFN-I and IFN-III locus in PH5CH8 hepatocytes; Related to** Figure 3**. (A-B)** Chromatin accessibility (ATAC-seq) and CUT&RUN signal for suppressive histone marks H3K9me3 and H3K27me3 in PH5CH8 hepatocyes at the IFN-I locus (chr9: 21.0-21.5 Mb) **(A)** and IFN-III locus (chr19: 39,225,486-39,326,550) **(B)** at 6 h.

**(C-D)** CUT&RUN signal for ATAC-seq, Pol II Ser 2P, H3K27ac, H3K4me3, H3K27me3 and H3K9me3 at individual IFN-I gene promoters **(C)** and IFN-III **(D)** gene promoters 1 kb upstream of TSS; gray dashed vertical lines indicate TSS position) in PH5CH8 hepatocytes at 6 h.

Mock (green) and RIG-ligand-stimulated (blue). Data in (A-D) are representative of one of 2 biological replicates.

**Supplementary Table 1**: HOMER motif enrichment analyses at IFN-I locus in pDCs and CAL-1 cells under mock and R848-treated conditions.

**Supplementary Table 2**: ETS-IRF composites motifs at PU.1 CUT&RUN binding peaks in CAL-1 cells under mock and R848-treated conditions.

**Supplementary Table 3**: Key resources table

## Acknowledgements

This project was in part funded by National Institutes of Health grants AI108765, AI135437, AR067980 (RS); T32AI165391 (MP); R35GM150503 (EAH). This project has been funded in whole or in part with Federal funds from the Frederick National Laboratory for Cancer Research, National Institutes of Health, under contract 75N91019D00024 (SA). This research was supported in part by the Intramural Research Program of NIH, Frederick National Lab, Center for Cancer Research. The contributions of the NIH authors are considered Works of the United States Government. The findings and conclusions presented in this paper are those of the authors and do not necessarily reflect the views of the NIH or the U.S. Department of Health and Human Services.

## Author contributions

Conceptualization: RS

Experimentation, data acquisition, data analysis: GM, SO, YO, MK, RS, HL, JS, APM, MP, EH, IMB-A, WL, SA Resource acquisition: RS, AL-H, JGS, EH, VN, TN, JAH, SA,

Writing – original draft: RS, GM

Writing, reviewing, and editing: GM, SO, YO, MK, RS, HL, JS, APM, DC, HW, MA, AL-H, JGS, MP, EH, IMB-A, WL, VN, TN, JAH, SA,

