## Supplementary Fig 1 for "Lineage-specific chromatin poising enforced by ETS-IRF composite elements determines the divergent interferon responses of plasmacytoid dendritic cells and epithelial cells"

**Supplementary Figure Legends:**

Supplementary Fig 1

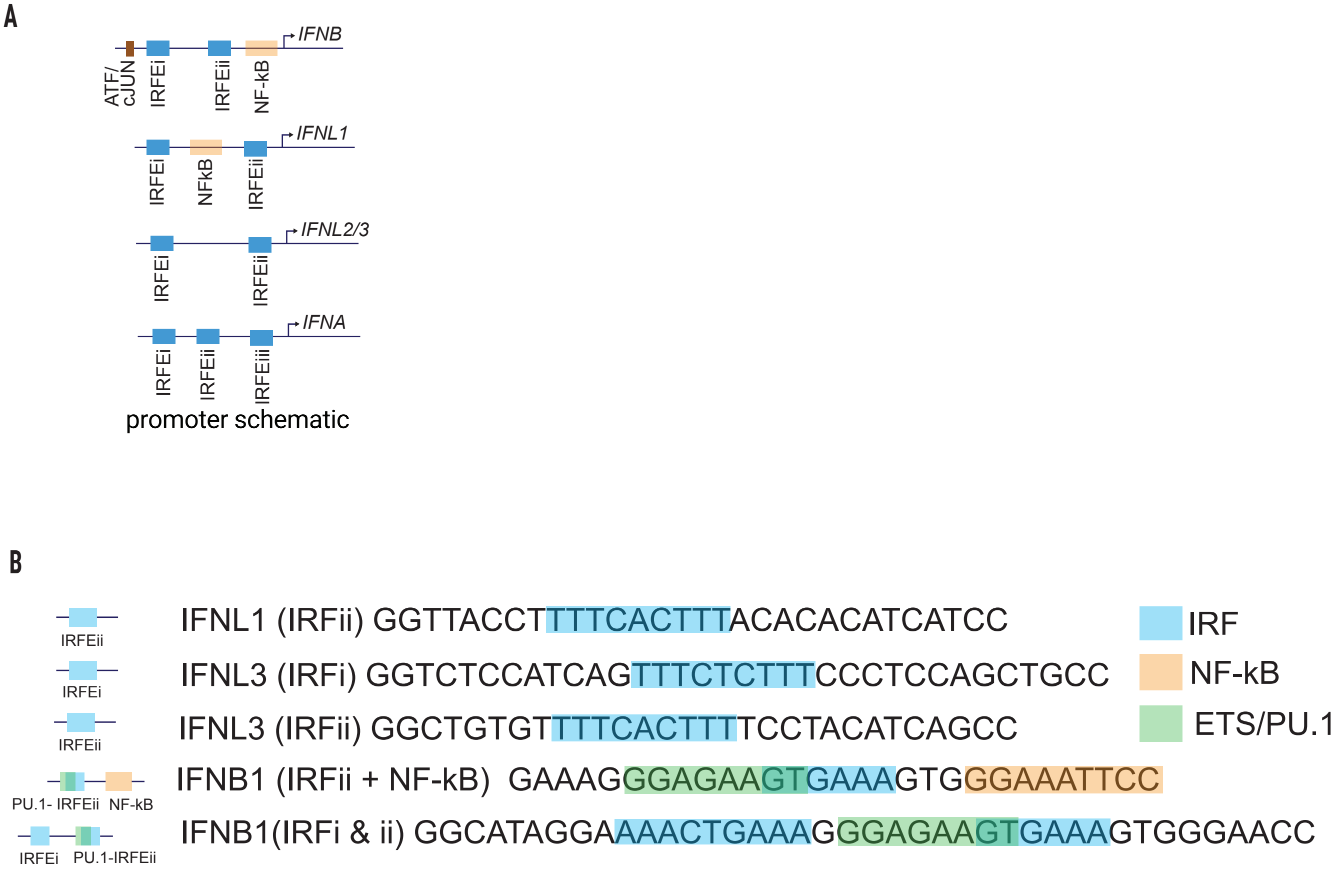


**Figure S1**: **(A)** Schematic of IFN promoters. **(B)**Schematic and genomics sequences of regulatory elements of IFNL1 IRFEii, IFNL3 IRFEi, IFNL3 IRFEii, IFNB1 (IRFEii+NF-κB), and IFNB1 (IRFEi+IRFEii). IRF binding sequences are shown in blue, NF-κB binding sequences in orange and ETS/PU.1 binding sequences in green. EMSA probes used experiments in Figure 5B are based on these promoter sequences.
