## Supplementary Fig 2 for "Lineage-specific chromatin poising enforced by ETS-IRF composite elements determines the divergent interferon responses of plasmacytoid dendritic cells and epithelial cells"

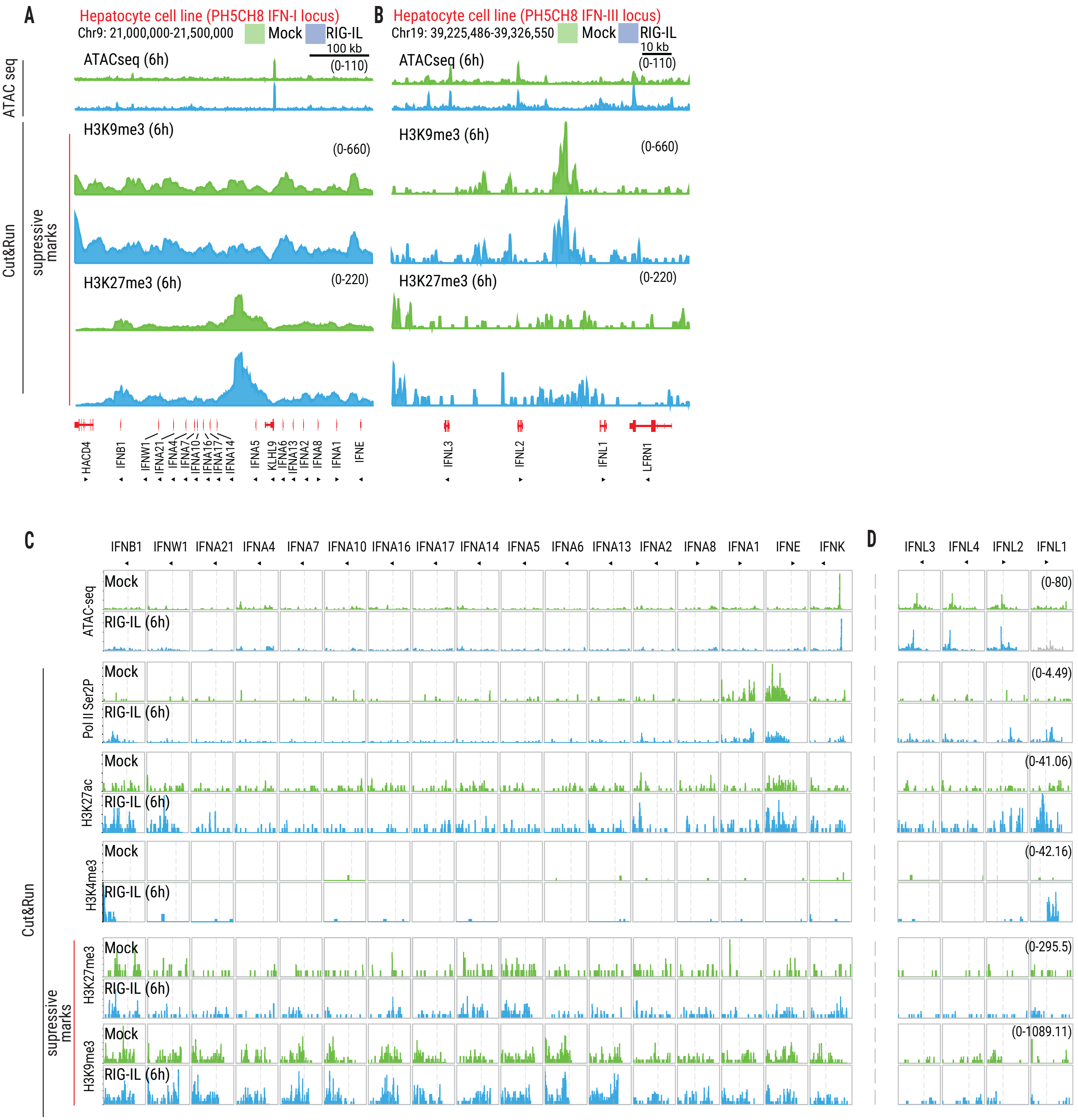


**Figure S2**: **Epigenetic landscape at IFN-I and IFN-III locus in PH5CH8 hepatocytes; Related to Figure 3.** **(A-B)** Chromatin accessibility (ATAC-seq) and CUT&RUN signal for suppressive histone marks H3K9me3 and H3K27me3 in PH5CH8 hepatocyes at the IFN-I locus (chr9: 21.0-21.5 Mb) **(A)** and IFN-III locus (chr19: 39,225,486-39,326,550) **(B)** at 6 h.

**(C-D)** CUT&RUN signal for ATAC-seq, Pol II Ser 2P, H3K27ac, H3K4me3, H3K27me3 and H3K9me3 at individual IFN-I gene promoters **(C)** and IFN-III **(D)** gene promoters 1 kb upstream of TSS; gray dashed vertical lines indicate TSS position) in PH5CH8 hepatocytes at 6 h.
Mock (green) and RIG-ligand-stimulated (blue). Data in (A-D) are representative of one of 2 biological replicates.
